# The dynamics of piRNA expression in *Blattella germanica* ovaries

**DOI:** 10.64898/2026.04.02.716027

**Authors:** D. Pujal, G. Ylla, J. Bau, M.D. Piulachs

**Affiliations:** Institute of Evolutionary Biology (CSIC-Universitat Pompeu Fabra). Barcelona. Spain; Department of Biosciences. University of Vic - Central University of Catalonia, Vic, Barcelona, Spain; Faculty of Biochemistry, Biophysics and Biotechnology. Jagiellonian University. Kraków, Poland

**Keywords:** cockroach, insect reproduction, sncRNA, insect oogenesis, panoistic ovary

## Abstract

The cockroach Blattella germanica possesses panoistic ovaries, in which oocytes lack nurse cells and therefore need to rely on their own transcriptional activity to support embryogenesis. Ovarian development in this species involves the development of a single basal ovarian follicle (BOF) per gonadotropic cycle, a process strictly regulated by endocrine signals, primarily juvenile hormone and ecdysone, which act at both the transcriptional and translational levels. In addition, transcriptional activity in these ovaries is necessary for both regulating and genome protection, and at this level, PIWI-interacting RNAs (piRNAs) play an essential role. Although insect ovaries are known to be particularly rich in piRNAs, their function in ovary maturation is still not well defined. For this purpose, we characterize the piRNA expression dynamics across seven key developmental and reproductive stages, ranging from late nymphal instars to post-vitellogenic adults.

piRNA expression in B. germanica shows coordinated fluctuations. Expression remains stable in previtellogenic ovaries, whereas vitellogenic ovaries show pronounced changes. Moreover, vitellogenic ovaries exhibit reduced piRNA diversity due to strong enrichment of a subset of highly expressed piRNAs.

Our data show that although piRNAs predominantly map to transposable elements, particularly LINEs, there is a notable increase in gene-derived piRNAs toward the end of the cycle. Our results suggest regulatory roles of piRNAs in modulating both TEs and mRNAs during BOF maturation, likely related to changes in the follicular cell program.

## Introduction

Ovarian development in insects is a complex and highly coordinated process regulated by molecular signaling cascades and hormonal control (1–3). Using *Blattella germanica* as a model, we investigate a new regulatory layer in insect oogenesis mediated by Piwi-interacting RNAs (piRNAs).

piRNAs are small non–coding RNAs (sncRNAs) initially identified in the male germline of *Drosophila melanogaster* (4,5). They have since been described in a wide range of eukaryotes, from plants to vertebrates. Their canonical function involves the regulation of transposable elements (TEs), from which they can also originate (5,6), thereby acting as essential genome guardians (7). However, accumulating evidence has revealed that the piRNA pathway extends beyond TE silencing. In insects, piRNAs participate in growth regulation, gonadal and wing development, embryo patterning, and sex determination (8–12). These findings show the importance of piRNAs as regulatory molecules, with some cases showing functional parallels to microRNAs.

In a previous study based on *B. germanica* development, we identified a total of 866,980 unique piRNAs across 11 developmental stages, from egg and embryo through successive nymphal instars to adult females (13). The abundance of piRNAs detected in unfertilized eggs suggests a significant role in ovarian physiology. These piRNAs appear to accumulate during oocyte growth, supporting ovarian maturation and potentially influencing early embryogenesis. Notably, *B. germanica* oogenesis is characterized by the growth of a single basal ovarian follicle (BOF) per ovariole in each gonadotropic cycle. Thus, most molecular changes observed in the ovary correspond to BOF development (reviewed in 2 and 3). This process is tightly regulated by juvenile hormone (JH) and ecdysone.

To elucidate piRNA function during oogenesis, we used small RNAseq to identify the piRNAs expressed in *B. germanica* ovaries throughout the gonadotropic cycle by sampling key developmental stages. These included ovaries with previtellogenic (immature), vitellogenic, and post-vitellogenic (mature) BOFs (Fig 1). We selected seven stages defined by hormonal peaks (JH or ecdysone) and by characteristic morphological changes in the follicular epithelium in the BOF (Fig 1).

**Fig 1.**
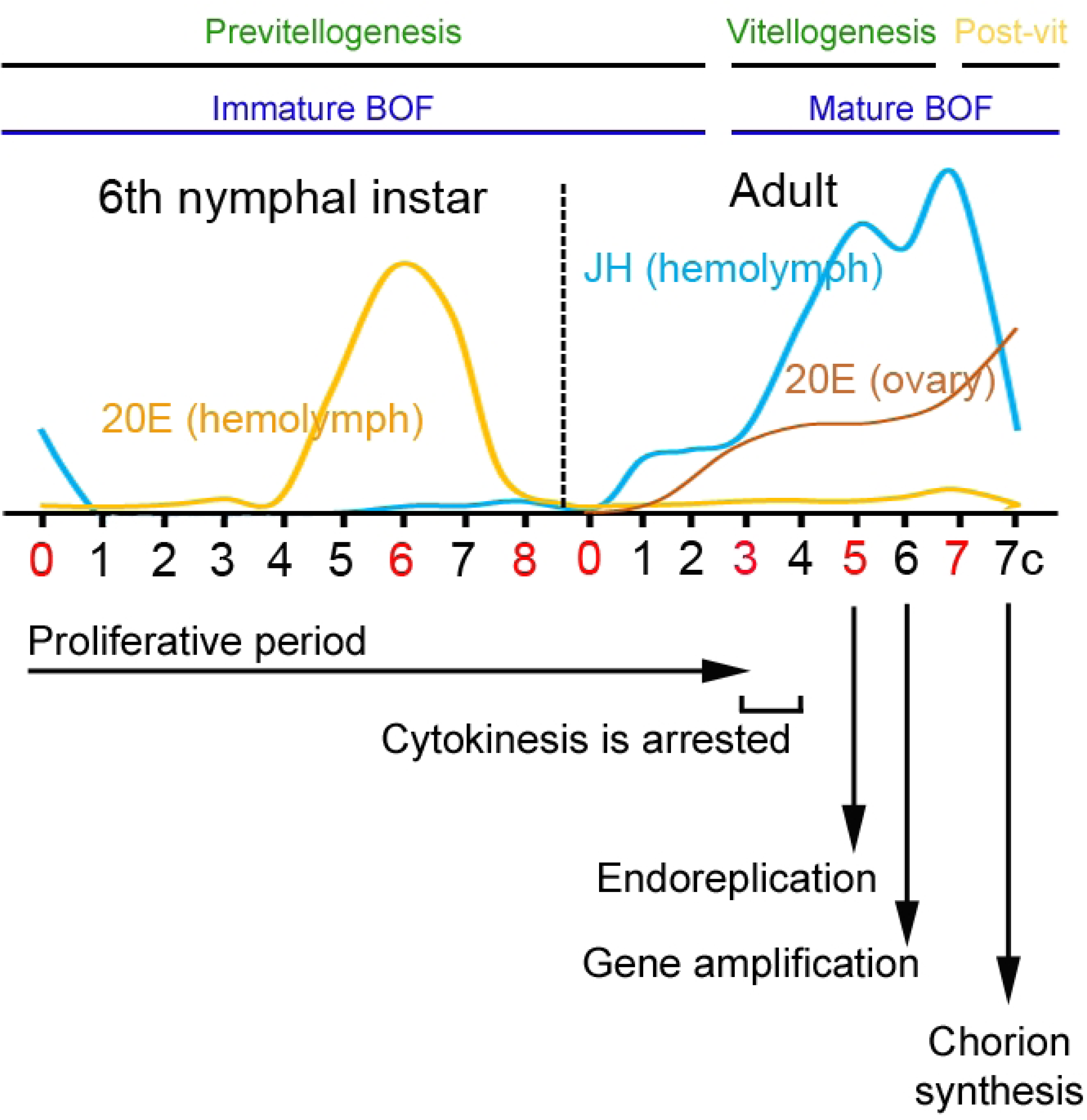
The oogenesis in *Blattella germanica*. Profiles of juvenile hormone (JH, blue) and 20-hydroxyecdysone (20Ec, yellow) in the hemolymph of last instar nymphs and adult females (7c indicates females with ovaries in choriogenesis) (data from Cruz et al., 2003; Pascual et al., 1992; Romaña et al., 1995). The profile of 20E content in the ovaries is represented in orange (data from Romaña et al., 1995). The days chosen for library construction, which coincide with significant changes in both hormones, are highlighted in red. Ovaries from newly emerged last instar nymphs (N6D0) with JH and no 20E, N6D6 has the highest levels of 20E in the hemolymph and the absence of JH, and N6D8 with low levels of both hormones, before the molt to the adult, were chosen. The adult ovaries included were: just after the molt (AD0), when hormone levels were very low. AD3, when JH levels in the hemolymph increase and induce vitellogenesis. AD5 in the vitellogenic period when JH in the hemolymph is at high levels, and the content of 20E in the ovary is increasing. AD7, just at the end of the cycle before the chorion formation, with the highest levels of JH in the hemolymph and the highest 20E content in the ovary. The progression of the follicular cell cycle in the BOFs is indicated below the plot, from the first day of the sixth instar nymph (N6D0) to chorion formation in the adult (AD7c).

Three stages corresponded to the last nymphal instar, when prothoracic-gland-derived ecdysone predominates in the hemolymph (14). During this instar, the basal oocyte becomes fully transcriptionally active and initiates the synthesis and organization of macromolecules required later for embryogenesis. The selected nymphal stages were: newly emerged last-instar nymphs, marking the onset of the first gonadotropic cycle (15); 6–day–old last-instar nymphs, when the number of ovarian follicles per ovariole is established, the germarium cell differentiation is arrested (16), and a hemolymph peak of 20–hydroxyecdysone (20E) occurs, activating the ecdysone signaling pathway (14,16,17); 8–day–old last-instar nymphs, just before the imaginal molt, when the basal oocyte has completed the accumulation of mRNAs and proteins required for embryogenesis (18).

After adult emergence, ovarian physiology is governed primarily by JH, which triggers the onset of vitellogenesis and BOF growth (2,19). In adults, ovarian piRNAs were analyzed at four stages: newly emerged adults, before the rise of JH in the hemolymph; 3–day–old adults, when the cytokinesis in BOF follicular cells arrests, ovarian ecdysone synthesis begins, and an increase in JH induces vitellogenesis in the fat-body; 5–day–old adults, when follicular cells become binucleated and polyploid (20) (15,20), JH levels peak (21), patency is established, and ovarian 20E increases to activate early chorion gene expression (16); 7–day–old adults, a post-vitellogenic stage in which choriogenesis is completed (20) just before ovulation and oviposition; JH levels decline (21), while ovarian 20E reaches its maximum (14).

Our results reveal a substantial proportion of ovarian piRNAs originating from genic regions. Their abundance varies during BOF maturation, increasing markedly in pre-oviposition stages while remaining relatively stable in immature ovaries. Together, these findings support a central role for piRNAs in regulating ovarian development and maturation in the cockroach *B. germanica*.

## Materials and methods

### Cockroach colony and sampling

Adult and last nymphal instar females of the cockroach *B. germanica* (L.) were obtained from a colony fed ad libitum on Panlab 125 dog chow and water, reared in the dark at 29 ± 1 °C and 60–70% relative humidity. Females were selected and used at appropriate ages: 1) newly emerged last instar nymphs (N6D0); 2) 6-day-old last instar nymphs (N6D6); 3) 8-day-old last instar nymphs (N6D8); 4) newly emerged adults (AD0); 5) 3-day-old adults (AD3); 6) 5-day-old adults (AD5); 7) 7-day-old adult females in choriogenesis (AD7). All dissections were performed on specimens anesthetized with carbon dioxide. Tissues were stored at -80 °C until use.

### RNA extraction and small RNA libraries preparation

Total RNA was isolated using the GenElute Mammalian Total RNA Kit (Sigma, Madrid, Spain). RNA quantity and quality were estimated by spectrophotometric absorption at 260/280 nm in a Nanodrop Spectrophotometer ND-1000® (NanoDrop Technologies, Wilmington, DE, USA). Small RNA libraries from *B. germanica* ovaries were prepared and sequenced by BMK (Münster, Germany) using the Illumina Novaseq 6000 (PE50, n = 3), except for AD3 samples (SE50, n = 4). Small RNA-seq data are publicly available at the NCBI Gene Expression Omnibus (22) under the accession code GEO: GSE255380. As a reference genome for the analyses, the *B. germanica* assembly generated by Harrison and colleagues (23) was used.

### Annotation of *Blattella germanica* transposable elements

*B. germanica* Transposable elements (TE) were identified using the EarlGrey TE annotation pipeline (v5.1.1; Docker image tobybaril/earlgrey_dfam3.7:latest). The Arthropoda repeat library from Dfam (v3.8) was used to mask known repeats, and TE sequences shorter than 100 bp were removed to generate the final repeat annotation. Gene models were adjusted to remove all overlaps with TE features using bedtools (v2.31.1) (24), and the resulting non-overlapping gene and TE annotations were used for downstream analyses.

### piRNA identification

Library quality was assessed using FastQC (v0.12.1) (25). Sequencing adapters were removed from the samples using Trimmomatic (v0.39, parameters: ILLUMINACLIP: TruSeq2-PE.fa: 2:30:10 LEADING:20 MINLEN:18) (26), and the trimmed libraries were subsequently subjected to another round of quality control with FastQC. Overlapping read pairs were merged using PANDAseq (v2.11) (27), and potential piRNA reads were then selected using Cutadapt (v3.5) (28), with a read length threshold set between 26 and 31 nucleotides. Reads were aligned to the *B. germanica* genome assembly (v1.1) (23) with Bowtie2 (v2.4.4, parameters: -k 1, --score-min L,0,0) (29). Only the reads that mapped with no mismatches were retained, while mapped reads containing one or more mismatches were removed using samtools (v1.19.2) (30) by selecting alignments with “NM = 0”.

To have a more consistent set of *B. germanica* piRNAs, reads that mapped to the same genomic locus and shared the first 26 nucleotides at the 5’ end, while only varying in their length at the 3’ end, were collapsed. The sequence with the highest expression among the sequences sharing the first 26 nucleotides was taken as the representative, and the expression of the representative and each 3’ variant was added to determine the total expression of the piRNA. Then, extremely infrequent piRNAs with an overall expression lower than 10 reads across all libraries were filtered out. The piRNAs obtained after this collapsing and filtering step were used in all subsequent analyses.

The identification of secondary piRNAs generated by the ping-pong amplification cycle involved searching for pairs of piRNAs with only 10 nucleotides of 5’-to-5’ perfect sequence complementarity. SeqLogos were generated with the R packages ggseqlogo (v0.2) (31) and ggplot2 (v3.5.1) (32).

To identify maternally derived piRNAs, two small RNA-seq libraries from non-fecundated eggs (NFE) were included ((33); GSE87031) and were processed using the same workflow as the other ovarian libraries.

### piRNA expression analysis

Read counts were normalized using the DESeq2 size-factor normalization method (DESeq2 v1.38.3) (34). The resulting DESeq2-normalized counts were used for downstream quantitative analyses, such as the application of expression thresholds. For visualization analyses, variance-stabilized transformed (VST) values were obtained using the varianceStabilizingTransformation function from DESeq2. VST values were used for Principal Component Analysis (PCA), the computation of Shannon’s diversity index (vegan v2.6-4), and for heatmap generation (pheatmap v1.0.12). Differential expression analysis between contiguous ovarian development stages was performed with DESeq2 (FDR < 0.05; log_2_FC > 2 | < -2).

### Characterization of piRNA genomic origins

The genomic origins of piRNAs were assessed by mapping the representative piRNA sequences to the *B. germanica* genome using Bowtie2 (v2.4.4, parameters: -a, --score-min L,0,0) (29), reporting all valid alignments for each sequence. Aligned reads with one or more mismatches were filtered out using samtools. The quantification of piRNAs mapped to genomic features (5’UTR, CDS, 3’UTR, introns, and transposable elements) was performed using the featureCounts function implemented in the R package Rsubread (v2.12.3, parameters: minOverlap=25, countMultiMappingReads=T, fraction=T) (35). Additionally, strandSpecific=1 was applied to count sense mappings and strandSpecific=2 for antisense mappings. The piRNA counts mapping to each genomic feature were normalized using TPM, accounting for the potential overrepresentation of piRNAs in longer genomic features.

Gene Ontology (GO) terms for *B. germanica* genes were obtained based on protein sequence orthology using eggNOG-mapper (version emapper-2.1.12) based on eggnog (36). Enrichment analysis of GO-terms was performed using Fisher’s exact hypergeometric test implemented in the R package topGO (v2.50.0) (37).

## Results

### The piRNA in the *Blattella germanica* ovary

Since piRNAs are primarily derived from genomic regions associated with TEs, we examined the *B. germanica* genome to determine the frequency of TEs in this species. Our annotation of the total repetitive content of the *B. germanica* genome identified 47.65% of the assembly as repetitive sequences. DNA transposons and LINEs are the most representative TE orders, spanning 10.93% and 9.71% of the genome, respectively (Table S1). The genome span of other TEs, such as LTRs, SINEs, and Rolling Circle, is much lower (0.49%, 0.45%, and 0.12%, respectively). Additionally, we identified other types of repeats, including simple repeats, microsatellites, low-complexity repeats, and unclassified repeats, which together account for 25.95% of the genome.

Here, we obtained 22 small RNA libraries from nymphal and adult *B. germanica* ovaries at different developmental stages, generating, on average, 13.13 ± 2.36 million reads per library. We selected reads ranging from 26 to 31 nucleotides, obtaining an average of 3.54 ± 0.80 million reads per library, of which 67.5% mapped to the reference genome (Table S2). After collapsing 3’ variants (identical sequences from the same genomic locus but differing at the 3’ end), we obtained 3,979,726 unique sequences. Due to the significant number of lowly expressed sequences, a filtering step was performed, removing sequences with fewer than 10 identical reads across samples, leaving a final set of 303,954 unique sequences that would represent the *B. germanica* ovarian piRNA repertoire.

*B. germanica* piRNAs (78.34%) are 28-29 nucleotides long, with 28-nucleotide piRNAs being the most common length (45.08%; Fig 2A). A smaller proportion of putative piRNAs are 27 nucleotides long (12.90%), while the rest corresponds to sequences of 26, 30, and 31 nucleotides (Fig 2A). As expected, the piRNA sequences display the typical U-bias at the first 5’ nucleotide (Fig 2B). A search for piRNAs involved in the ping-pong pathway, namely with a 10-nucleotide overlap at their 5’ end and mostly with an A in position 10 (Fig 2D), revealed 84,369 piRNA sequences forming 274,606 overlaps, representing 27.8% of all the identified piRNAs, most of them with a length of 28-29 nucleotides (Fig 2C).

**Fig 2.**
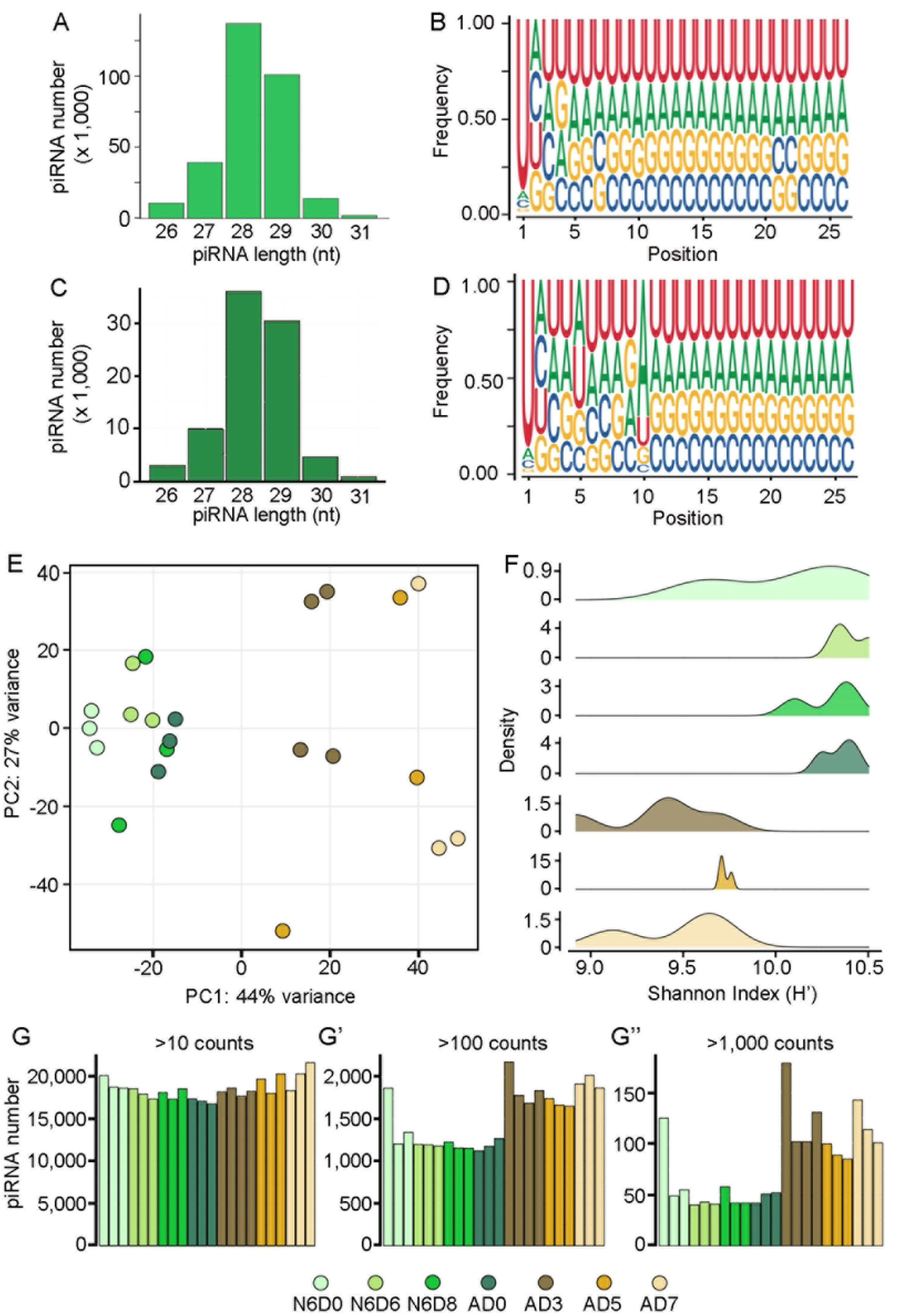
piRNAs identified in the *Blattella germanica* ovary. **A.** Number of piRNAs distributed by length (nt: nucleotides). **B.** Sequence logo for all identified piRNAs showing the nucleotide frequency in each position, with a clear bias for an uracil (U) in the first 5’ position. **C.** Distribution of *B. germanica* secondary pathway piRNAs (defined by a 5′ 10-nt overlap with another piRNA) by length (nt: nucleotides). **D.** Sequence logo displaying the nucleotide frequency of piRNAs from the secondary pathway, showing the uracil (U) and adenine (A) biases in the 1^st^ and 10^th^ positions, respectively. **E.** Principal component analysis showing the distribution of ovarian piRNA libraries of last instar nymphs and adults. **F.** Density plot depicting the Shannon’s diversity index (H’) distribution of ovarian piRNA stage-libraries. **G.** Number of piRNAs with expression over 10 counts, (G’) 100 counts, and (G’’) 1,000 counts distributed by stage in each library. N6 indicates last instar nymphs and A the adults. The letter D, followed by a number, indicates the age of the individual in days within the instar.

### piRNA expression dynamics in the *B. germanica* ovary

The dynamics of piRNA expression in the ovaries, during the first gonadotropic cycle, were analyzed using PCA (Fig 2E). The analysis revealed a clear divergence of samples along the developmental timeline, reflecting substantial changes during BOF development (Fig 2E). The first principal component (PC1) accounted for 44% of the variance, while the second (PC2) explained 27%. Ovaries of the last nymphal instar and newly emerged adults (AD0) formed a tight cluster, indicating that piRNA expression remains highly similar along those stages. Within this cluster of previtellogenic ovaries, the earliest stage (N6D0) is slightly separated along the first principal component (PC1), suggesting a minor shift in piRNA expression following the emergence to the last nymphal instar. At this moment, the BOF oocytes are previtellogenic, undergoing capacitation to enter a vitellogenic stage, while the follicular epithelia exhibit a high degree of proliferation (Fig 1).

In contrast, samples of AD3, AD5, and AD7 ovaries show a broader dispersion along the PC1, indicating substantial changes in piRNA expression over oocyte maturation (Fig 2E). The PCA analysis revealed a clear difference between ovaries from previtellogenic (N6D0 to AD0) and vitellogenic (AD3 to AD7) stages, considering piRNA content and expression. These differences appear to be especially marked between immature and AD3 ovaries, as well as between AD3 and AD5 ovaries (Fig 2E). The dispersion along PC2 evidenced variability across replicates in vitellogenic ovaries, which is possibly explained by the physiological transitions in the BOF associated with the crucial changes in the follicular cells program that take place over short time intervals at the end of the gonadotropic cycle.

The analysis of piRNA diversity, using Shannon’s diversity index (H’), revealed two distinct profiles (Fig 2F), which correspond to the clustering observed in the PCA. Libraries from immature ovaries exhibited greater diversity of piRNAs, as reflected by higher H’ values (Fig 2F), suggesting that the expression in these stages is more evenly distributed across different piRNAs. In contrast, vitellogenic ovaries exhibited reduced piRNA diversity, suggesting that during these stages, a smaller subset of distinct piRNAs is expressed at higher levels (Fig 2F).

We examined piRNA abundances across a range of expression thresholds to clarify how the expression of several piRNAs contributes to the patterns seen in the PCA and Shannon’s diversity analysis (Figs 2E and 2F). The number of piRNAs with expression levels above 10 DESeq2-normalized counts is consistent across all ovarian libraries (Fig 2G). However, when the threshold is raised to 100 DESeq2-normalized counts, a clear distinction emerges between mature and immature ovaries (Fig 2G’). The highest expressed piRNAs are more abundant in mature ovaries. When we further extend the threshold to piRNAs expressed above 1,000 DESeq2-normalized counts, the difference between mature and immature ovaries becomes even more evident, with mature ovaries containing about twice the amount of piRNAs compared to immature stages (Fig 2G’’).

Differential expression analysis between consecutive stages of ovarian development, revealed significant variation across these stages (Figs 3 and S1). In the first transition, from N6D0 to N6D6, only 76 piRNAs were differentially expressed, with 30 being upregulated and 46 downregulated in ovaries from N6D6 (see Supplementary Fig S1).

**Fig 3.**
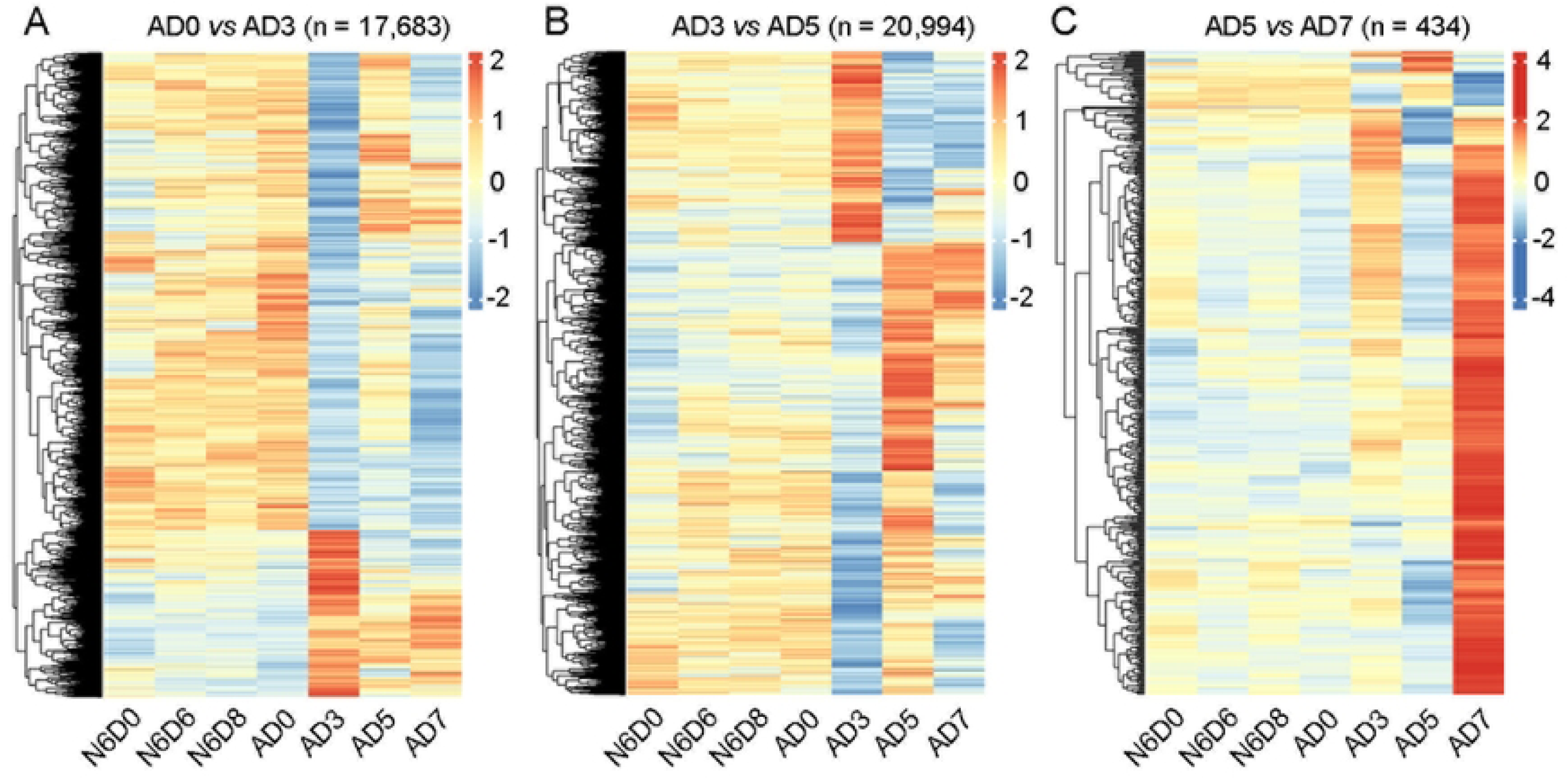
Differential expression of *Blattella germanica* piRNAs throughout ovary maturation. Heatmap Plots showing the piRNAs differentially expressed in the transitions: 0-day-old adult (AD0) to AD3 **(A)**, AD3 to AD5 **(B)**, and AD5 to AD7 **(C)**; with a log_2_FC > 2 | <-2 and a False Discovery Rate (FDR) < 0.05. Expression of the differentially expressed piRNAs is depicted for all stages (N6D0 to AD7) to show their expression profile throughout the whole ovary maturation process. In the transition N6D0 to N6D6, only 76 piRNAs were found to be differentially expressed (see Supplementary Fig S1), whereas the transitions N6D6 to N6D8 and N6D8 to AD0 resulted in 0 and 1 DE piRNAs, respectively.

Most of these piRNAs maintain their expression in immature ovaries. Interestingly, the subsequent transitions (N6D6 to N6D8 and N6D8 to AD0) showed very small changes in piRNA expression, with only 0 and 1 differentially expressed piRNAs, respectively. All of these suggest the presence of a stable set of piRNAs in immature ovaries, likely essential for maintaining cellular functions, particularly in the BOF.

The transition from an immature to a mature BOF (AD0 to AD3) is critical in the *B. germanica* ovary. In AD3, the proliferation of follicular cells in the BOFs begins to arrest, while oocytes grow by accumulating storage proteins, mainly vitellogenin (Fig 1). During this transition, a significant number of piRNAs (17,683) are differentially expressed (Fig 3A). One major group is downregulated in AD3 as the BOFs begin to mature; subsequently, some of these piRNAs increase in expression again as the BOFs mature, approaching the end of the gonadotropic cycle. In contrast, a second group of piRNAs expressed at lower levels in previtellogenic ovaries shows elevated expression in AD3, maintaining these levels until the end of the gonadotropic cycle.

In *B. germanica* females, during the transition from AD3 to AD5, JH levels in the hemolymph increase, and 20E begins to accumulate in the ovary. The follicular cells in the BOF arrest cytokinesis and become binucleated, entering polyploidy (Fig 1). During this transition, a notable change in piRNA expression occurs, with 20,994 piRNAs identified as differentially expressed (Fig 3B). Many of the piRNAs appear upregulated in AD5, and mostly maintain their expression levels in AD7, suggesting a specific role in the later stages of the gonadotropic cycle. Conversely, a group of piRNAs is downregulated in AD5. These downregulated piRNAs maintain the low expression levels until oviposition.

The final transition, from AD5 to AD7, is critical for completing the gonadotropic cycle. In the ovaries of 7-day-old females, follicle cells in the BOF undergo endoreplication in specific genomic regions, leading to upregulation of chorion genes (20). During this transition (Fig 3C), a total of 434 piRNAs are differentially expressed, exhibiting a wide range of fold changes. Notably, 91.5% of these piRNAs are upregulated, with an average fold change of 3.9, ranging from 2.0 to 22.6 (Fig 3C). Their expression levels are higher than at any other age. In contrast, at AD7, only a limited number of piRNAs are downregulated; 37 piRNAs show a decrease in expression when compared to AD5, with fold changes ranging from 2.0 to 6.3 (Fig 3C).

The significant increase in upregulated piRNAs in AD7 (Fig 3C) raises an important question regarding their function: Are these piRNAs acting within the ovary, or are they being transferred to the egg? To address this question, we utilized the small RNA libraries corresponding to eggs from virgin females (NFE: non-fecunded eggs;13,33), where we identified 10,749 piRNAs exclusive to NFE, indicating that they likely originate from the mature oocyte and thus represent piRNAs from the germinal line (Fig S2A). These maternal piRNAs represent 9.8% of the total piRNAs present in NFE and are found generally at low read counts (Fig S2B). The rest of the piRNAs identified in NFE (98,041 piRNAs), which are also present in mature ovaries, are present in greater proportions (Fig S2C). These data suggest that the piRNAs maternally deposited in the egg participate in regulating some functional processes in early embryo development.

### The genomic origins of piRNAs

To better understand the piRNA functions in *B. germanica*, we investigated the genomic regions where these piRNAs mapped without any mismatches. To ensure valid comparisons between different features and samples, the number of mapped locations was normalized as previously described (see Section 2.6). The adjusted values were then used to calculate the mapping proportion of piRNAs to each feature within each sample.

As expected, the main sources of piRNAs correspond to TEs (Fig 4A), with 88.9% of the ovarian piRNAs mapping to TE sequences. We analyzed which TEs piRNAs map to and whether they align to the sense or antisense strand of the TE (Figs 4A and S3). Among Class I TEs, piRNAs are in a relatively stable density throughout all stages of ovarian development (Figs 4A and S3), except for those piRNAs mapping SINEs in the sense strand, which are less abundant in ovaries at the end of the gonadotropic cycle, becoming significantly reduced in ovaries from AD7, just before oviposition. Between this Class I, we found that the largest number of piRNAs mapped to LINEs, in both sense and antisense strands (Figs 4A and S3). Conversely, among Class II TEs, the proportion of piRNAs mapping to DNA TEs increases significantly at the end of the cycle, compared to the rest of the days (Fig S3), whether they map on the sense or the antisense strand. The piRNAs mapping the antisense strand of RC decrease significantly in the last days of the gonadotropic cycle (Figs 4A and S3).

**Fig 4.**
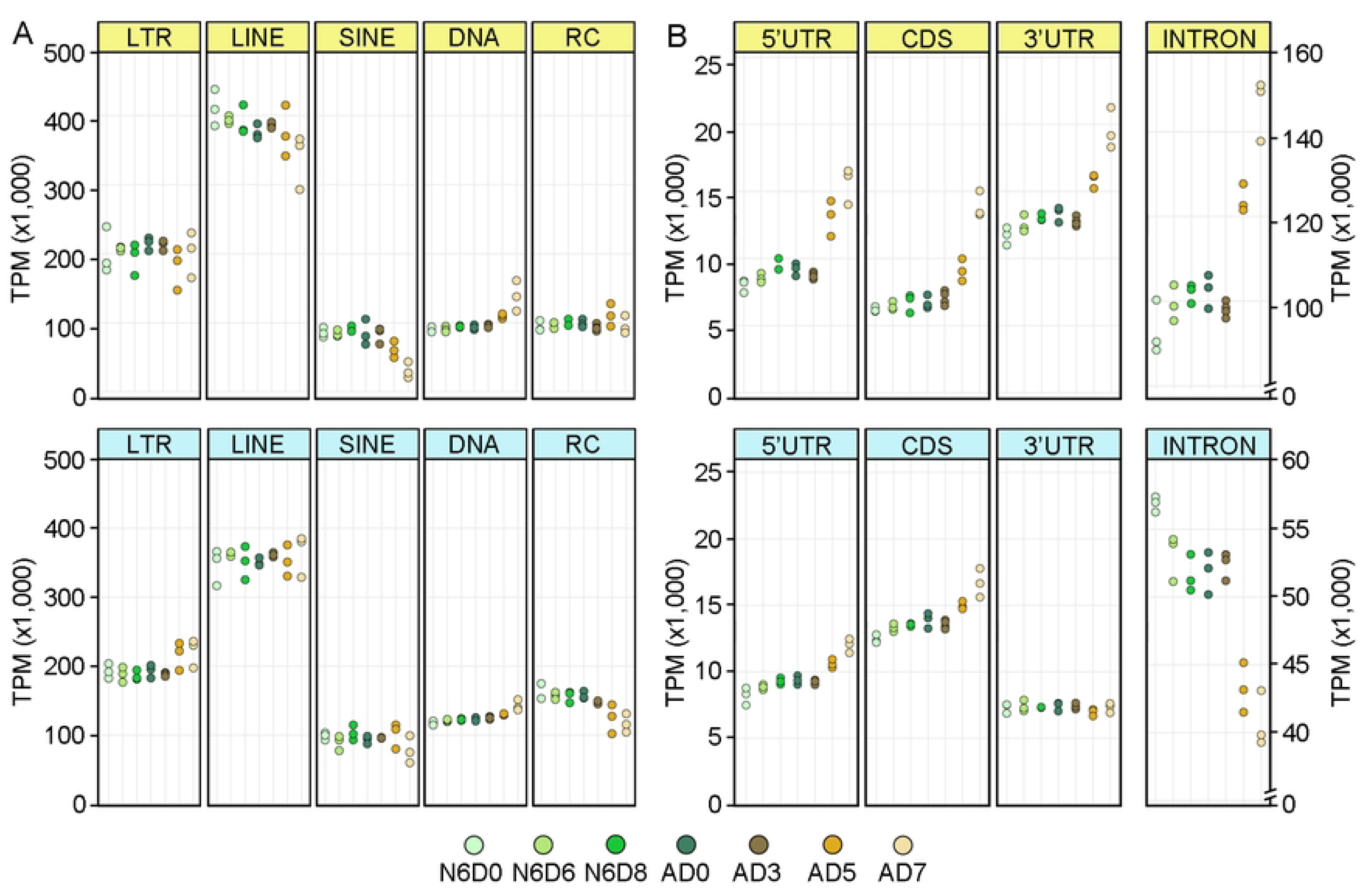
Genomic mapping sites of ovarian piRNAs in *Blattella germanica*. **A.** TPM of piRNAs mapping to the five main TE orders (LTR, LINE, SINE, DNA, and RC) in the sense strand. **B.** TPM of piRNAs mapping to the genic regions (5’UTR, CDS, 3’UTR, and Introns) in the sense strand. **C.** TPM of piRNAs mapping to the five main TE orders (LTR, LINE, SINE, DNA, and RC) in the antisense strand. **D.** TPM of piRNAs mapping to the genic regions (5’UTR, CDS, 3’UTR, and Introns) in the antisense strand. N6 indicates last instar nymphs, and A the adults. The letter D, followed by a number, indicates the age of the individual in days within the instar.

In addition to TEs, a significant portion (11.1%) of piRNAs map to genic regions (3.2% in UTRs and CDS, 7.9% in introns), in both strands (Figs 4B and S4). Showing significant changes across the different stages as the BOF matures (Fig 4B and S4). In the later stages of the gonadotropic cycle, in AD5 and AD7, there is a significant increase in piRNAs mapping to CDS and 5’UTRs (Fig S4) in both strands. Changes that also occur in those piRNAs mapping the sense strand of 3’UTRs, suggesting an increase in regulatory functions for piRNAs in the ovaries at these ages. The high number of piRNAs mapping introns is stable in immature ovaries. However, in ovaries from AD5 and AD7, a significant increase of piRNAs in the sense strands occurs, while in the antisense strand, with a low number, a clear decrease of piRNAs is observed (Figs 4B and S4).

The proportion of piRNAs mapping to genic regions led us to explore which type of genes correspond to these piRNAs in the *B. germanica* genome. In total, we identified 4,026 gene where piRNA maps, from either strand (sense and antisense): 66.8% in CDS, 11.3% in 5’UTR, and 21.9% in 3’UTR. A Gene Ontology (GO) enrichment analysis of these genes identified GO annotations for the Biological Process (BP) category in 803 genes. The most significantly enriched GO BP terms were mainly related to metabolic processes, DNA conformational changes, transport, reproduction, gene expression, and ribosome biogenesis (Fig S5).

Additionally, when analyzing the piRNAs mapping to genic regions such as CDS, 5’UTR, and 3’UTRs, we identified a small group of piRNA pairs that align to opposite strands in the same genomic locus with a 10-nucleotide difference in their start positions, similar to those described by Jensen and collaborators (38). The analysis found a total of 18 genes presenting intrinsic ping-pong pairs, whose roles cover various biological processes, including autophagy and lysosome positioning (3/18), chromatin regulation (3/18), ribosome biogenesis (1/18), transcription regulation (1/18), and transposon-derived proteins (4/18), while six genes encode for proteins with unknown functions.

Taken together, our data demonstrate that piRNA expression undergoes coordinated changes during the transition from previtellogenic to vitellogenic stages, suggesting that these piRNAs may play essential roles at the end of the gonadotropic cycle to complete BOF development.

## Discussion

The piRNAs are a class of molecules with gene regulatory properties on TEs and RNAs (mRNAs and long non-coding RNAs) (39). In arthropods, piRNA expression has been found in various gonadal and somatic tissues (40–42). Among these, the ovaries, containing both somatic and germinal cell lineages, exhibit the highest abundance of piRNAs (43,44).

TEs represent the primary genomic locations where piRNAs originate, TEs represent the primary genomic locations where piRNAs originate, and Petersen et al. (45) and Ylla et al. (46), suggest that TE occupancy generally increases with genome size in insects, although notable variability across species has also been documented. In the wasp *Nasonia oneida*, 32.29% of the genome consists of repetitive sequences (47), while in the honey bee *Apis mellifera*, only 9.46% of the genome is covered by repetitive sequences, while TEs represent only 3% (48). In Diptera, the percentage of TEs also varies significantly, from 20% in the *Drosophila melanogaster* genome (49) to 47.7% of TEs in *Aedes aegypti* (50). This range is also present in hemimetabola insects. Among Orthoptera, in the cricket *Gryllus bimaculatus*, the TEs represent 28.94% of its genome (46), while in the locust *Locusta migratoria*, the presence of TEs reaches 65% (45). Our analysis determined that in the cockroach *B. germanica*, 47.65% of the genome is occupied by repetitive sequences, and among them, 21.7% are annotated TEs, with DNA TEs being the most abundant type (10.93%). Contrasting with the predominant presence of LINE in the Gryllidae, that is 20.21% of annotated TEs in *Laupala kohalensis* (46). In other species, like *D. melanogaster* and *Tribolium castaneum*, LTR retrotransposons are the most abundant class of TEs (51,52). Remarkably, SINEs, although present in all insect orders, are less abundant (45).

Given the abundance of TEs in insect genomes, it is expected to find a very high number of piRNAs in insects. This is the case of *R. prolixus* (53) and *Aedes albopictus* (54), with about 36 million and 12 million piRNAs reported, respectively. These numbers, however, may be overestimated, suggesting that more stringent criteria should be applied to accurately identify biologically active piRNAs and enable studies of their potential regulatory roles (55). Applying stringent conditions in *B. germanica*, collapsing identical sequences with varying levels of trimming and mapping at the same genome locus, and selecting the piRNA sequence with the highest expression, we significantly reduced the number of piRNAs previously identified (13). With the resulting 303,954 *B. germanica* piRNAs, it was possible to analyze their dynamic expression in ovaries. The PCA analysis provides initial evidence of the potential role of piRNAs in *B. germanica* ovarian maturation, showing clear distances between previtellogenic and vitellogenic ovaries, which could be determined by piRNA origin and number, at each stage.

We found that the different TE families contribute similarly to the ovarian piRNA pool. Although the number of piRNAs mapping to some TE families is maintained throughout the gonadotropic cycle, it is in the last days of the cycle when it varies substantially. This is the case of piRNAs mapping the sense strand of SINA or the antisense strand of RC, which reduces their number significantly. Conversely, piRNAs mapping on DNA TEs, sense or antisense, increase significantly on the last day of the cycle. This data suggests that TEs play a role at the end of the cycle, either by providing piRNAs or being depleted by them.

The piRNAs mapping to genic regions (CDS and UTRs) in *B. germanica* are approximately 3.5%, a relatively low proportion compared to *L. migratoria*, another species with panoistic ovaries, where 11% of piRNAs are located in genic regions, including 8% in CDS (41). In other species with meroistic ovaries, such as the bug *Oncopeltus fasciatus* and the bumblebee *Bombus terrestris*, the proportion of piRNAs mapping to CDS is notably higher (25% and 48%, respectively), while the proportion mapping to UTRs remains relatively constant (41).

In *B. germanica*, we observed significant changes in the abundance of piRNAs mapping to genic regions when comparing different stages of ovarian development. Notably, at the end of the gonadotropic cycle, in adult females, 5- and 7-day-old, we detected the greatest variability in the abundance of piRNAs mapping to the 5’ UTR and CDS and introns, in both the sense and the antisense strand. While for those piRNAs mapping the 3’UTR, the differences at the end of the cycle correspond to the piRNAs mapping the sense strand, while the ones mapping the antisense strand remain stable.

All this data suggests a role of the piRNAs participating in the changes observed in the follicular cell program that occurs in the basal ovarian follicles (BOF) at the end of the gonadotropic cycle (Fig 1), where big changes occur at the genome level, to facilitate the chorion synthesis (3,20).

Analyzing the piRNA expression across the different stages of *B. germanica* ovarian development, a few differentially expressed piRNAs (76 piRNAs) were detected in immature ovaries (N6D0-AD0), where piRNA expression is very stable. However, some piRNAs modify their expression just after the molt to the last nymphal instar, showing a significant downregulation. These expression changes could be related to the entrance of the *B. germanica* ovary in the maturing period, which starts early in this last nymphal instar and is marked primarily by the stabilization of the number of ovarian follicles in the ovarioles and the initiation of BOF maturation (3,16). Thus, the piRNA content in *B. germanica* ovary, and their expression levels clearly categorize the previtellogenic ovaries (with immature BOF), including those of newly molted adults, and separate them from mature ovaries (Fig 2E). In mature ovaries, the changes in piRNA expression align with the most significant cell changes in the follicular epithelium in the BOF. In AD5, there is a set of piRNAs downregulated, suggesting that their functions could be related to the changes in the program of the follicular cells, which become binucleate and polyploid (3,15). In AD7, the last day of the gonadotropic cycle, a small set of piRNAs is differentially expressed, with some of them reaching high expression levels.

In summary, our study reveals that the ovarian piRNA landscape of *B. germanica* is dynamic, with stable periods and sharp transitions that could be aligned with major endocrine and cellular events of the gonadotropic cycle. Furthermore, we set up the basis for future avenues to investigate how small RNAs integrate hormonal signalling, follicular cell dynamics, and gene regulation in the ovary of hemimetabolous insects.

## Supporting Information

**S1 Fig. Differential expression analysis of piRNAs in ovaries of *Blattella germanica* in the transition N6D0 to N6D6.** The heatmap plot depicts the expression profile of differentially expressed piRNAs throughout the ovarian maturation, with a log2FC > 2 | < -2 and a False Discovery Rate (FDR) < 0.05.

**S2 Fig. Maternal transmission of piRNAs in *Blattella germanica***. A. Comparison of piRNA content in the mature ovarian stages (AD3, AD5, and AD7) and non-fecunded egg (NFE). The UpSet plot was generated with the R package ComplexHeatmap (v2.26.0) using the “distinct” mode for combination size calculation. B. Number of NFE-exclusive piRNAs at different count levels. C. Number of piRNAs shared between mature ovarian libraries and NFE libraries at different count levels. The count levels in B and C, were calculated as the log10 of the mean of the DESeq2 normalized counts in NFE (n = 2).

**S3 Fig. Mapping sites of Blattella germanica ovarian piRNAs, in annotated TEs**. piRNAs mapping the different TE orders: LTR. LINE, SINE, DNA, RC, in sense and antisense strands, expressed in TPM. Data is presented as mean ± S.E.M. (n = 3, except AD3 n = 4). The asterisks indicate statistically significant changes: * = p< 0.03, ** = p< 0.01, *** = p< 0.001, **** = p< 0.0001, ns= no significant.

**S4 Fig. Mapping sites of Blattella germanica ovarian piRNAs, in genic features**. piRNAs mapping genic regions: 5’UTR, CDS, 3’UTR and Introns, in sense and antisense strands, expressed in TPM. Data is presented as mean ± S.E.M. (n = 3, except AD3 n = 4). The asterisks indicate statistically significant changes: * = p< 0.03, ** = p< 0.01, *** = p< 0.001, **** = p< 0.0001, ns= no significant.

**S5 Fig. Gene Ontology enrichment analysis of genes with piRNA origins in their sequence.** 4,026 genes were identified and selected for the analysis. The GO annotations for Biological Process were retrieved for 803 of these genes. Gene Ratios were calculated as the number of genes annotated with a given GO Term, divided by the total number of input genes. Enriched GO Terms were ordered by increasing p-values, and the top 20 terms were represented.

## Notes

### Competing Interest Statement

The authors have declared no competing interest.

